# PEI-Coated Microbubble Attachment to Neutrophils: Potential for Radiation Force assisted delivery into tissue

**DOI:** 10.64898/2025.12.15.693345

**Authors:** Hossein Razmi Bagtash, Roshni Gandhi, Ghazal Rastegar, Amine Azizi, Aparna Priyadarshani Jha, Shuai Shao, Emma Salari, Shashank R. Sirsi, Caroline N. Jones

**Author notes:** These authors contributed equally to this work (Hossein Razmi Bagtash, Roshni Gandhi).

## Abstract

Immunotherapies have advanced cancer treatment; however, their clinical efficacy remains limited for solid tumors due to challenges associated with effectively directing immune cells into the complex tumor microenvironment. Recent developments in Ultrasound Contrast Agent (UCA — also known as “microbubble”) technology have provided novel opportunities to enhance targeted therapeutic delivery. In this study, we introduce an innovative approach of leveraging microbubbles to enhance immune cell targeting by directly attaching microbubbles to immune cells, enabling the targeted delivery and localization of immune cells into solid tumors using radiation force ultrasound (US) application. To create novel microbubble-immune cell conjugates, we created polyethyleneimine (PEI) coated microbubbles and attached them to differentiated HL-60 (dHL-60) cells. These positively charged PEI microbubbles were formulated using azide-DBCO click chemistry between DBCO-labeled microbubbles and the azide functional groups on the PEI polymer. Following this step, we utilized electrostatic interactions to attach our positively charged PEI microbubbles to our negatively charged dHL-60 cells. We conducted viability experiments to assess the compatibility of these designs and verified that cell viability remained greater than 88% four hours after the conjugation process for different ratios of dHL-60 cells to PEI microbubbles. We used microfluidic chemotaxis platforms to quantify the microbubble-conjugated dHL-60 cell migratory behavior, examining parameters including migration velocity and percentage. Additionally, we investigated the impact of ultrasound power on primary human neutrophils to validate the functional responsiveness of these physiologically relevant immune cells. Here we demonstrated the possibility of ultrasound-responsive immune cell constructs as a targeted strategy without loss of function in migration capabilities. The novel PEI microbubble and immune cell conjugates reported in this work will be used to improve future immunotherapy techniques.

## Introduction

The tumor microenvironment (TME) plays a pivotal role in determining the efficacy of cancer immunotherapies(1). Its complex and dynamic landscape establishes diverse physical, chemical, and immunological barriers that obstruct the effective infiltration and activity of immune cells within tumor tissues (1,2). Specifically, the pronounced heterogeneity of the TME, its dense extracellular matrix (ECM), and abnormal vasculature impede immune cell migration and limit their cytotoxic capabilities upon reaching the tumor core (3,4). In addition to these structural challenges, the TME exerts active immunosuppressive effects through factors such as hypoxia, acidosis, and the accumulation of immunosuppressive cell types, including regulatory T cells and myeloid-derived suppressor cells (1,3). To overcome this limitation, strategies that deliver pre-activated or pre-treated immune cells directly into the tumor hold promise for enhancing their functional activity and persistence in these hostile environments (1,2,5–7). Cancer immunotherapy relies on using the natural killing mechanisms of the immune system to eliminate tumor cells (2,8) by either activating an antigen on the tumor or enhancing the immune system itself. Given the current limitations of immunotherapy, especially in solid tumors, a method of delivering immune cells to the complex tumor microenvironment can help improve efficacy of this therapy. Many immunotherapy strategies aim to enhance immune cell infiltration into tumors by manipulating chemokines that attract effector cells to the tumor microenvironment (4,9). Directly introducing specific subpopulations of immune cells into solid tumors, rather than relying on tumor-driven immune cell recruitment, has the potential to revolutionize the clinical application of immunotherapy. However, instead of conventional intratumoral injection approaches, employing focused ultrasound (FUS) for targeted delivery offers several distinct advantages(10). First, ultrasound enables broader spatial coverage within the tumor and surrounding stroma, ensuring more uniform modulation and minimizing regions that remain inaccessible to injected cells(11–13). Second, it reduces procedural risks associated with needle-based delivery, such as bleeding, infection, tumor seeding, and patient discomfort, making it safer and less invasive, particularly for deep or delicate anatomical sites(14,15). Third, FUS provides noninvasive access to multifocal or anatomically challenging lesions, guided by real-time imaging for high precision(10,16). Together, these advantages position focused ultrasound as a powerful alternative for enhancing the safety, efficacy, and translational potential of intratumoral immunotherapy.

Microbubbles are composed of an exterior shell that can be made up of lipids, polymers, or proteins that encapsulate a gas core(17). These microbubbles are similar in size to red blood cells and stay in circulation for *in vivo* applications, they can be employed for a variety of therapeutic purposes(17). Moreover, they are acoustically responsive and can move in the direction of a propagating ultrasound wave; this phenomenon is known as primary radiation force application(18). This unique aspect of microbubbles makes them versatile cargo delivery vehicles, meaning therapeutic drugs and genes can be delivered to specific areas of the body(17,19–21). Cells can be delivered using microbubbles for *in vitro* and *in vivo* applications as a way to improve upon immunotherapy(22–24). Previous studies have demonstrated the ability to conjugate Sonazoid microbubbles to natural killer cells for cell therapy and imaging(25). Other studies have demonstrated the ability to deliver stem cells using microbubbles to treat post myocardial infarction atherosclerosis(23). Further research is needed to explore different types of microbubble and cell conjugates, identify optimal ratios of these compounds, and translate them into clinical trials for immunotherapy. We have developed a novel microbubble and cell conjugate for immunotherapy purposes using polyethyleneimine (PEI) microbubbles and neutrophils. By pre-conjugating our microbubbles to immune cells, we propose to enable their site-specific delivery to the solid tumor using low-intensity radiation force ultrasound. As a crucial initial step, it is necessary to show that the microbubble conjugation process onto immune cells does not affect their function or activity, which is the main focus of this study.

To evaluate the effects of microbubble conjugation on immune cell viablility and function, a microfluidic platform to monitor cell migration was used. Such platforms enable the quantification of immune cell migration function in physiologically appropriate, controlled settings(26–29). Microfluidics technologies have been used to quantify immune cell-function(26,30–33). These platforms have been used to define dynamic processes including chemotaxis(34,35), cell-cell interactions(36,37), and immunological responses(30,36–39). Microfluidic systems can measure immune cell migration in linear gradients of chemoattractants(32,40,41).

In this study, a microfluidic system was utilized to explore the migratory patterns of immune cells conjugated with microbubbles, allowing for real-time observation of how microbubble conjugation and ultrasound power influences cell migration to chemotactic signals. Neutrophil-like differentiated HL-60 (dHL-60) cells have been used in this study, as HL-60 cells are frequently used as model systems to investigate neutrophil function. DHL-60 cells exhibit characteristics of neutrophils, including chemotaxis, bactericidal activity, neutrophil extracellular trap (NET) creation, and reactive oxygen species (ROS) generation (42,43). dHL-60 cells provide a reproducible, genetically tractable, and scalable system for studying neutrophil behavior, including migration, all of which are critical to our microbubble-conjugation and ultrasound-based targeting strategy (44,45). To validate our findings in a physiologically relevant context, we also performed key functional experiments using primary human neutrophils. These primary cells were used for migration studies under ultrasound stimulation. Importantly, the results observed in primary neutrophils were consistent with those obtained using dHL-60 cells, supporting the relevance of the cell line as a model system while ensuring that our conclusions are applicable to actual human immune cells. This approach not only supports the development of targeted delivery strategies but also deepens our understanding of immune cell function within engineered microenvironments. We validate the formulation of a novel type of polyethyleneimine (PEI) microbubbles which is a branched cationic polymer that has the ability to carry more cargo due to its chemical structure(46). Moreover, because of the click chemistry, PEI microbubbles are fast and stable(47). We confirmed that the attachment of PEI microbubbles to immune cells does not have a detrimental impact on viability and migration.

## MATERIALS AND METHODS

### dHL-60 cell culture

HL-60 cells (ATCC CCL-240) were purchased from the American Type Culture Collection (ATCC, Rockville, MD, USA). Cells were cultured in Iscove’s Modified Dulbecco’s Medium (ATCC) and supplemented with 20% fetal bovine serum. Cells were cultured at 37°C and at 5% CO_2_. In order to bring the cells to a differentiated state, dimethyl sulfoxide (DMSO, Sigma-Aldrich, St. Louis, MO) was added and the cells were incubated for 4-5 days. dHL-60 cells were used for preliminary viability and conjugation experiments due to their reproducibility and ease of culture.

### Primary Human Neutrophils

Five milliliters of peripheral blood were collected from healthy donors into heparinized tubes. Human neutrophils were isolated using the Human Neutrophil Isolation Kit (Stemcell Technologies) following the manufacturer’s protocol. After isolation, the cells were adjusted to the desired concentration and mixed with PEI-coated microbubbles at specific ratios. Following exposure to defined ultrasound power, the neutrophils were injected into the microfluidic system to assess chemotaxis and migratory behavior over a 4-hour period using time-lapse imaging. Primary neutrophils were used for migration studies to ensure physiologically relevant functional responses.

### Preparation of microbubbles

Lipid microbubbles for PEI conjugation were prepared using a similar protocol that has been described previously(48). Notably, in this study we have changed the conjugation chemistry from maleimide-thiol to copper free Strain-Promoted Azide-Alkyne Cycloaddition (SPAAC) click chemistry. Lipid films were prepared with 1,2-distearoyl-sn-glycero-3-phosphocholine (DSPC), N-(methylpolyoxyethyleneoxycarbonyl)-1,2-distearoyl-sn-glycero-3-phosphoethanolamin (DSPE-PEG2000), and 1,2-distearoyl-sn-glycero-3-phosphoethanolamine-N-poly (ethylene glycol)-dibenzocycolctyne (DSPE-PEG-DBCO) using a molar ratio of 90:5:5 respectively. Each lipid was dissolved in a chloroform (Sigma-Aldrich, St. Louis, MO) stock solution and mixed into a single 20 mL glass vial to achieve the desired lipid ratios. This mixture was nitrogen evaporated and stored at -20 °C. The stored 5% DBCO film was suspended in 10 mL of 1X phosphate buffer saline (PBS, Fisher Scientific, Waltham, MA), 10% propane-1,2,3-triol (glycerol, 92.09 FW) (v/v), and 10% propane-1,2-diol (propylene glycol, 76.1 FW) (v/v), creating a 2 mg/mL solution. An Isotemp Heating Block was used to heat the solution up to 65 °C. In order to completely suspend the lipid, the solution was bath sonicated using an Ultrasonic Bath Sonicator (Fisher Scientific, Waltham, MA). A probe tip sonicator with a microtip attachment at maximum amplitude for 10 seconds was used to emulsify the lipid solution with Decafluorobutane (PFB, 238 MW, FluoroMed LP, Round Rock, TX) gas to form the microbubble suspension. The microbubbles are then cooled in an ice bath and centrifuged three times using a Bucket Centrifuge Model 5804R (Eppendorf, Hauppauge, NY) at 300 relative centrifugal force (RCF) for 3 min to separate the microbubbles from free lipid. The size distribution and concentration of the final bubble suspension was measured by a Multisizer 4e Coulter Counter (MS4, Beckman Coulter, Brea, CA).

### Azide-PEI conjugation to DBCO-Microbubbles

Azide-labeled branch polyethylene glycol polyethylenimine (N3-PEI-PEG) with a PEI molecular weight of 10k, PEG molecular weight of 2k, and a 20% substitution ratio (Nanosoft Polymers, Winston-Salem, NC) was conjugated to the microbubbuble surface using a protocol similar to our previous work (49). The suspension volume for 10 mg/mL of PEI-g-PEG is determined using the molecular weight and size distribution data of the 5% DBCO microbubbles. Then, 100 uL of microbubbles are slowly mixed in the solution consisting of PEI-g-PEG and 1XPBS. The microbubbles and PEI-g-PEG solution were stored in a 3 mL Luer tip syringe. This solution is then mixed thoroughly for 60 minutes using a Laboratory Tube Rotator to allow sufficient time for the azide-DBCO covalent bonds to form. This solution is then washed three times using the same settings described above for the centrifuge and characterized again using the Coulter Counter.

### PEI microbubble conjugated dHL-60 cell preparation

The HL-60 cells (CCL-240, ATCC) were mixed with PEI microbubbles, shown in **Figure 1A**, along with the microbubble generation process, at ratios of 1:1,1:2, 1:5, and 1:10. These ratios were calculated using the concentration and volume of the HL-60 cells and PEI microbubbles. Schematic design of the formulation of PEI microbubbles using 5% DBCO microbubbles and polyethyleneimine (PEI). Due to electrostatic interactions, the negatively charged HL-60 cells will bind to the positively charged PEI microbubbles **(Figure 1B)**.

**Figure 1.**
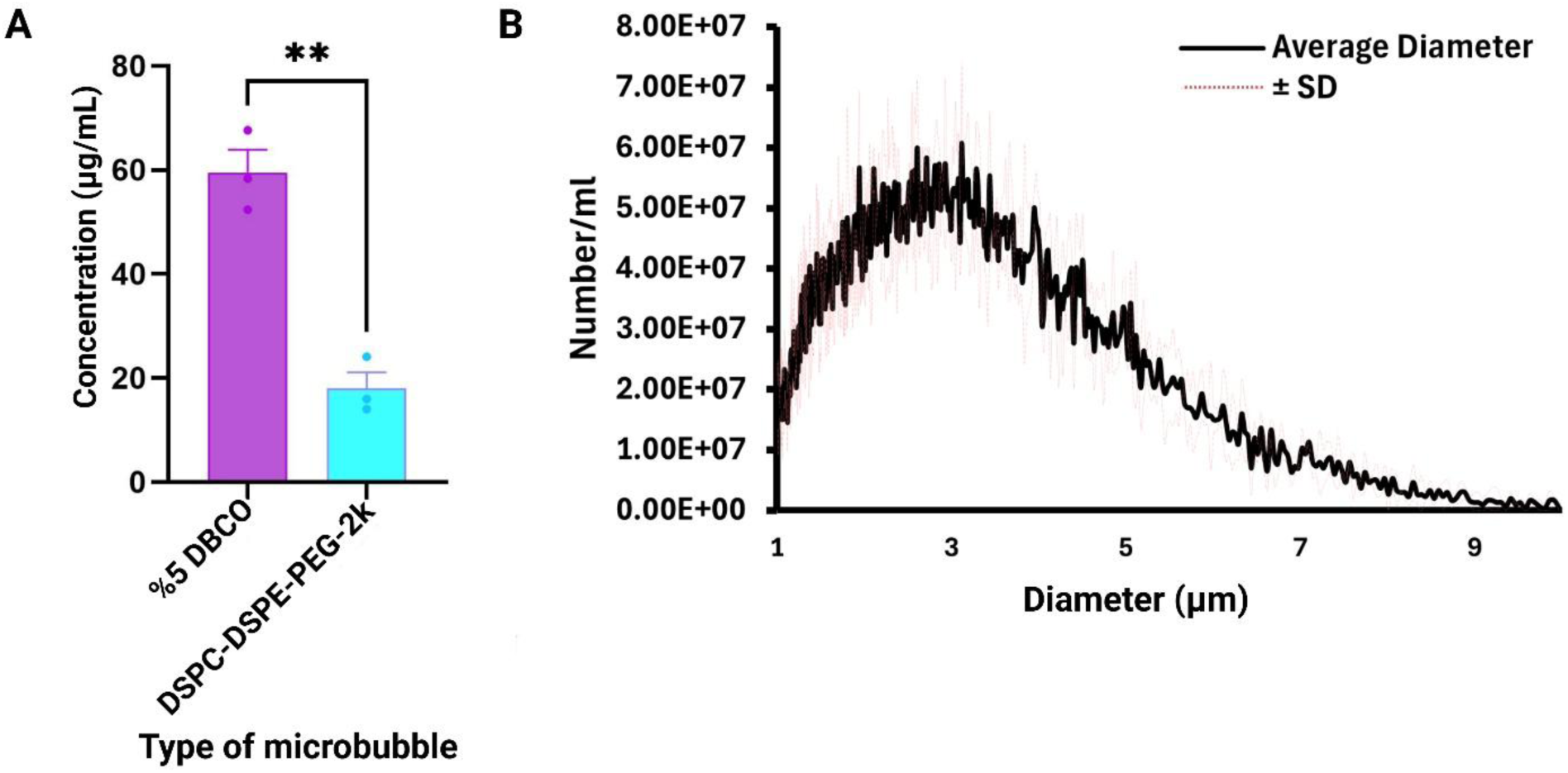
Preparation and conjugation process of PEI microbubbles with HL-60 cells. **(A)** Schematic design of the formulation of PEI microbubbles using 5% DBCO microbubbles and polyethyleneimine (PEI). **(B)** PEI microbubble conjugated HL-60 cells were made by utilizing the electrostatic forces between the positively charged PEI microbubbles and negatively charged HL-60 neutrophils. Created with BioRender.com.

### Fluorescamine detection assay

Dilutions of 10 ug/mL to 1000 ug/mL were made using polyethyleneimine (PEI) (Sigma-Aldrich, St. Louis, MO) and phosphate buffered saline (PBS). A fluorescamine stock solution with a concentration of 3 mg/mL was made (Sigma-Aldrich, St. Louis, MO) with dimethyl sulfoxide (DMSO) (Sigma-Aldrich, St. Louis, MO). Aliquots of 90 µL of the polymer stocks were pipetted into a 96 well microplate. PEI microbubbles and DSPC-DSPE microbubbles (acting as a negative control) were also pipetted into the microplate and diluted with 1xPBS to reach a final volume of 90 µL and concentration of 3E9 #/mL. Each well was mixed with 30 µL of fluorescamine. The fluorescence was detected using a fluorescence plate reader with an emission filter of 460 nm and an excitation filter of 400 nm.

### Live/dead assay of dHL-60 cells

After treatment with microbubbles, dHL-60 cells were seeded on a glass-bottom 96-well plate (Cellvis, P961.5HN) at a concentration of 8 × 10^5^ cells/mL in 100 µL of complete IMDM. To assess the effect of microbubble treatment on the viability of dHL-60 cells, dHL-60 cells were stained with the LIVE/DEADTM Viability/Cytotoxicity Assay Kit (Invitrogen, L32250) based on the manufacturer’s protocol. Briefly, a 2X dye solution was prepared in complete IMDM and 100 µL of the dye solution was added to each well to reach a working concentration of 1X at t=0 h and 4 h after microbubble treatment. Cells were imaged on a Nikon ECLIPSE Ti2-E microscope using a Plan Apo 20X objective (NA = 0.80) at 37°C to capture live and dead cells at t=0 h and t=4 h. Images were acquired using NIS-elements (Nikon Inc.) software and recorded using FITC (live cells) and Cy5 (dead cells) channels. Cells were stored in the incubator at 37°C with 5% CO2 between the two time points of imaging. Image analysis was performed in ImageJ. Cell viability was quantified as the percentage of live cells per image, i.e., the number of live cells divided by the sum of the number of live cells and the number of dead cells. The number of cells was measured using Thresholding and Analyze Particles functions. The Watershed function was used to segment clumps of cells. Only objects with a larger area than 15 µm^2^ were classified as cells to exclude spurious spots and debris.

### PEI microbubble formulation

PEI microbubbles were created using azide-DBCO click chemistry. A fluorescamine assay was used to detect the amine groups present in the azide-DBCO reaction. We determined that more PEI was coated on the 5% DBCO microbubbles than the DSPC-DSPE-PEG2k microbubbles since the 5% DBCO microbubbles had an average concentration of 59.5 µg/mL while the DSPC-DSPE-PEG2k microbubbles only had an average concentration of 18.05 µg/mL (**Figure 2A**). The DSPC-DSPE-PEG2k microbubbles did not have any DBCO amine groups present, and therefore a click chemistry reaction did not occur; any residual PEI left over after the centrifugation steps could have led to the values detected for the control group. Size distribution of PEI microbubbles was normal with a final concentration of 8.5E9 #/mL and mean diameter of 2.568 µm (**Figure 2B**).

**Figure 2.**
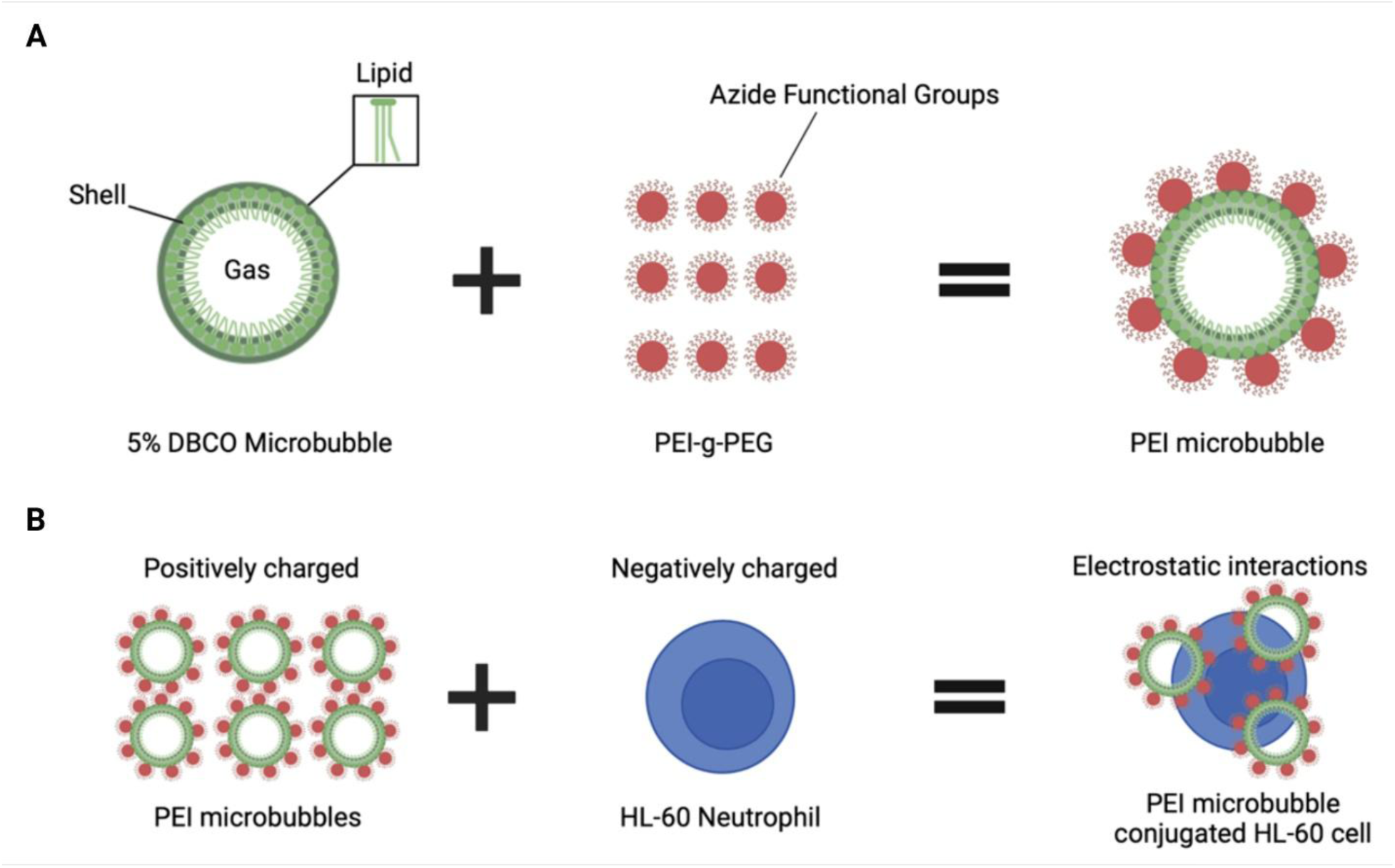
Characterization of PEI microbubbles. **(A)** PEI microbubbles were synthesized using azide-DBCO click chemistry. Results show that there is a greater concentration of PEI detected using the fluorescamine assay for the 5% DBCO microbubbles than DSPC-DSPE-PEG-2k microbubbles since more amine groups were detected after the click-chemistry reaction. **(B)** Size distribution of PEI microbubbles from the Coulter Counter. The PEI microbubbles have an average concentration of 8.532E9 microbubbles per mL and a mean diameter of 2.568 µm. Data are presented as mean ± SD from three independent experiments. Statistical significance was assessed using an unpaired two-tailed Student’s t-test. **p < 0.01.

### Microfluidic device preparation along with chemotactic migration of dHL-60 cells conjugated with PEI microbubbles

Microfluidic devices were fabricated using traditional photolithography techniques. Silicon mold was used to create devices with two chemoattractant reservoirs for generating linear gradients toward a central reservoir, where dHL-60 cells were cultured to study their migratory behavior. Each device also included ten migration channels (10 µm width × 10 µm height) connecting the central reservoir to the left and right chemoattractant reservoirs (**Figures 3A, B**). This process also utilizes replication molding, in which a PDMS prepolymer is created by mixing PDMS and a curing agent. The PDMS prepolymer is poured into the silicon mold of desired structure; it is then cured and later punched with a biopsy puncher. Oxygen plasma bonding (Harrick Plasma) was used to attach a glass slide to the bottom of the device. The migration patterns of the HL-60 cells were determined using microfluidic chemotaxis assays. Fibronectin, a glycoprotein in the extracellular matrix, was used to coat the microfluidic channels in order to increase cell adhesion. The PEI microbubbles and HL-60 cells were mixed at ratios of 1:1, 1:2, 1:5, and 1:10. These conditions were loaded onto the dHL-60 cells chamber of multiple devices **(Figure 3A)**. Chemoattractants N-Formylmethionine-leucyl-phenylalanine (fMLP, Sigma-Aldrich, St. Louis, MO) and Leukotriene B4 (LTB4, Cayman Chemical, Ann Arbor, MI) were loaded into two chemoattractant chambers at a concentration of 100 nM. Time lapse images were taken using the Nikon TiE fully automated microscope (Nikon Inc., Melville NY). Images were recorded at 2:30 minute intervals for 4 hours. The number of cells migrating towards fMLP and LTB4 and the number of PEI microbubble conjugated HL-60 cells were quantified. **Figure 3B** shows a real-time microscopic image of the microfluidic system, highlighting dHL-60 cells within the cell loading chamber and migration channels.

**Figure 3.**
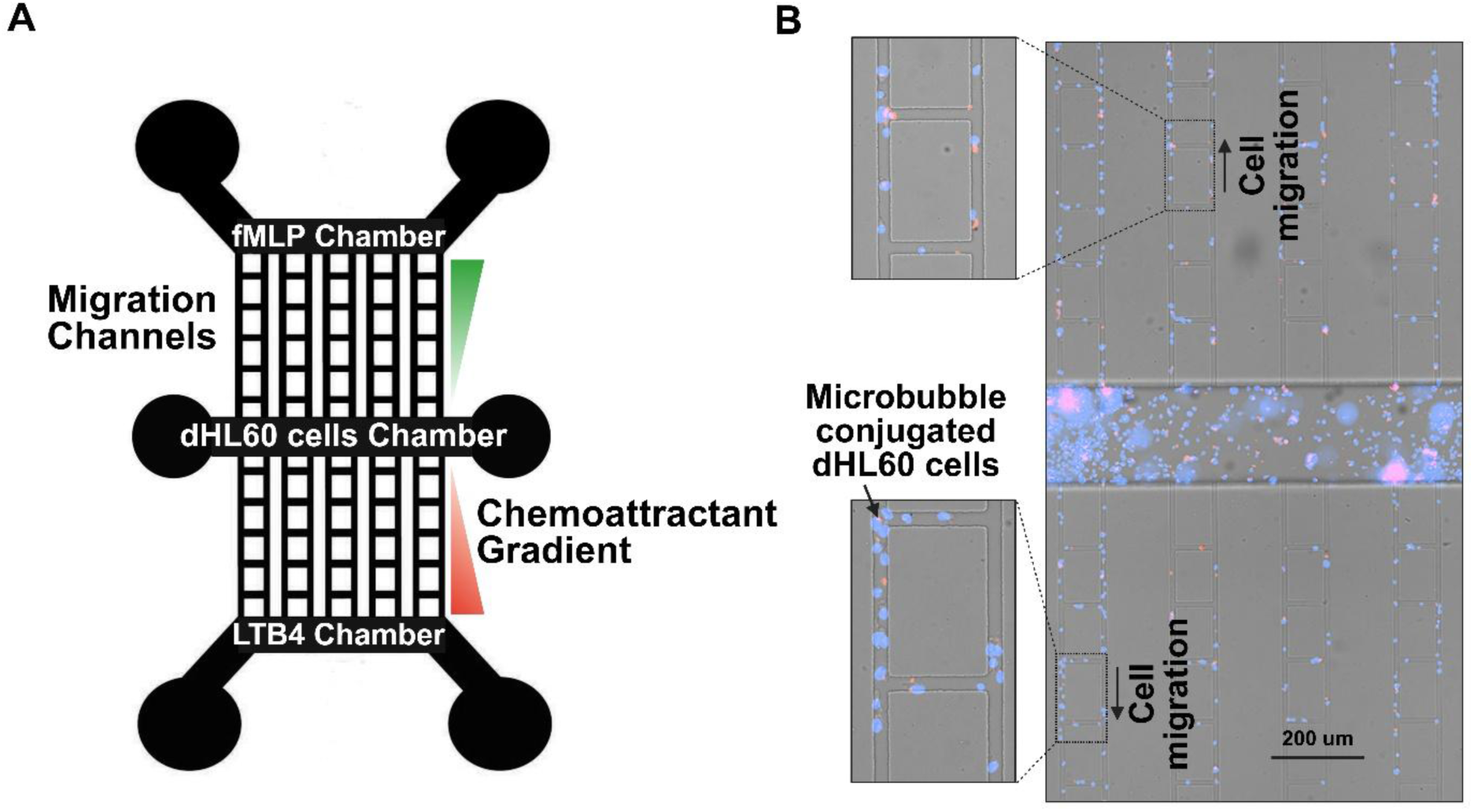
Design of a microfluidic device, chemotactic movement of dHL-60 cells conjugated with PEI microbubbles. **(A)** The microfluidic device’s schematic illustration. An array of migration channels (10 µm width, 10 µm height) connects the device’s central reservoir to two chemoattractant reservoirs (fMLP and LTB4). Cells migrate along the channels in response to chemoattractant gradients created by the fMLP and LTB4 reservoirs, while the central reservoir acts as the loading point for cells. **(B)** Microscopic image of the microfluidic system showing PEI microbubble-conjugated dHL-60 cells migrating toward the fMLP and LTB4 chemoattractant reservoirs in the upper and lower chambers. 200 µm is represented by the scale bar.

### Ultrasound Treatment of Microbubble-Conjugated Neutrophils

To evaluate the effect of ultrasound power on primary neutrophil migratory behavior, microbubble-conjugated neutrophils were exposed to ultrasound at different power levels prior to microfluidic migration assays. While most experiments in this study were conducted using differentiated HL-60 (dHL-60) cells, human primary neutrophils were employed in this section to validate the observed ultrasound effects in physiologically relevant cells.

Ultrasound exposure was performed using the Siemens Sequoia ultrasound with the 15L8 Linear transducer (Dr. Lux’s laboratory, UTSW Medical Center). Cells were treated under four conditions: (1) control (no ultrasound), (2) low-power ultrasound, (3) medium-power ultrasound, and (4) high-power ultrasound. The specific acoustic power settings for the low, medium, and high-power conditions were -30 dB, -10 dB, and 0 dB, respectively. Supplementary Video 1 demonstrates the visual response of microbubble-conjugated cells to 0 dB ultrasound application. The ultrasound acoustic radiation force was done using continuous pulse wave doppler mode to emit radiation force at different outputs.

For each condition, neutrophils were conjugated to microbubbles at a 1:2 cell-to-microbubble ratio. The cell–microbubble suspensions were transferred to sterile Eppendorf tubes and subjected to ultrasound exposure for 10 seconds at room temperature. Following ultrasound treatment, the cells were immediately introduced into the microfluidic migration assay platform.

Neutrophil migration was quantified as described in the migration assay section. The percentage of migrating cells was compared across all ultrasound conditions to assess the impact of ultrasound power on neutrophil motility.

### Ethics Statement

Primary human neutrophils were isolated from peripheral blood obtained from healthy adult donors under a protocol approved by the Institutional Review Board (IRB) at the University of Texas at Dallas. All donors provided written informed consent prior to participation.

## RESULTS

### Microbubble treatment does not impact the viability of dHL-60 cells in 4 h

We investigated the viability to determine the effect of PEI microbubble conjugation process on the survival of dHL-60s (**Figure 4A, B**). Previous literature states that microbubbles are stable, specifically polymer-based microbubbles(17). The viability of the PEI conjugated dHL-60 cells was compared with the control dHL-60 cells. A LIVE/DEAD^TM^ Viability/Cytotoxicity assay was used to determine the viable and nonviable cell count, and the percent viability. The percent viability of the PEI microbubble conjugated HL-60 cells was compared to that of the control dHL-60 cells. There was no significant difference between the percentage viability of the PEI microbubble conjugated dHL-60 cells and that of the control dHL-60 cells which have been shown in **Figure 4A, B**.

**Figure 4.**
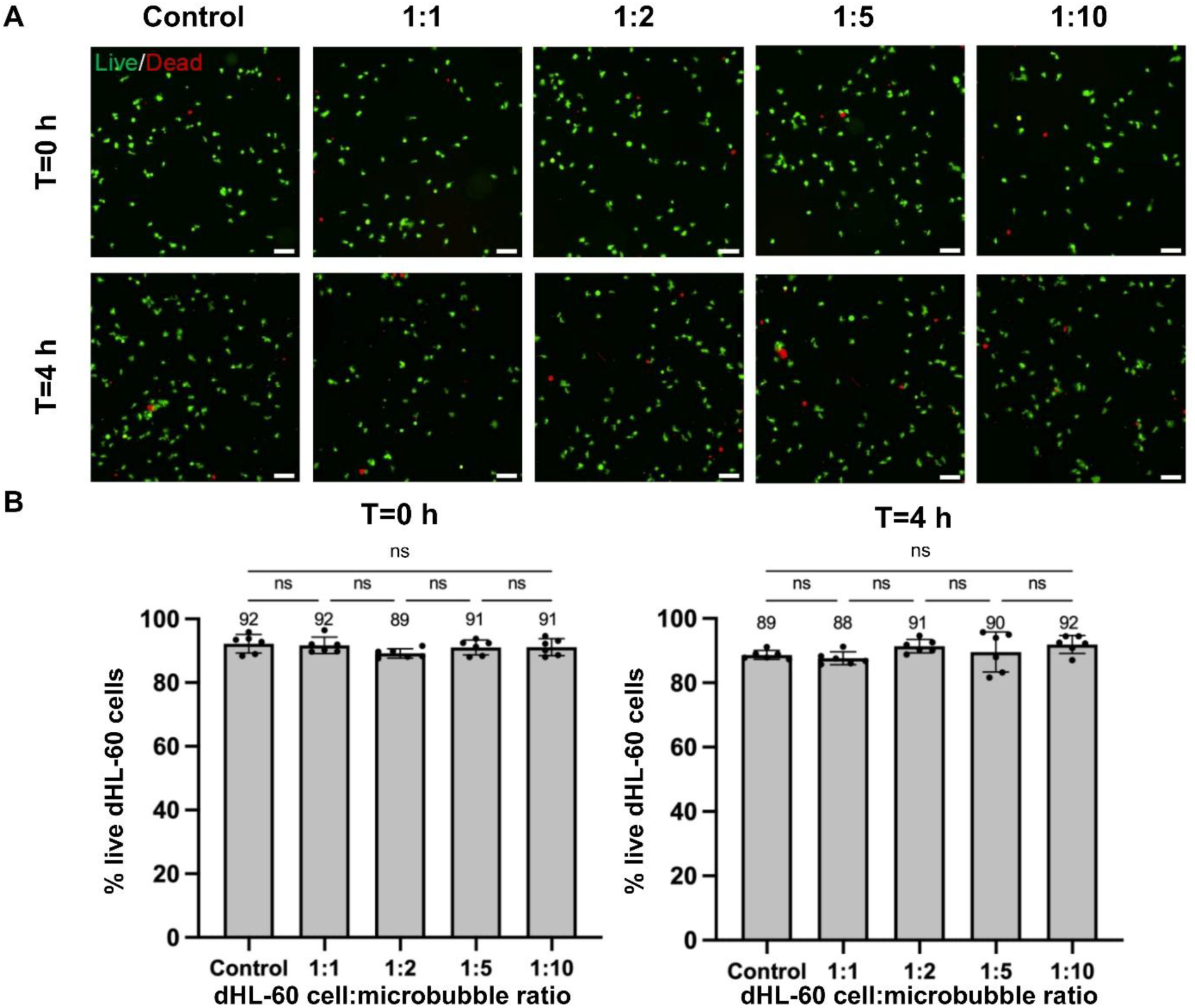
Microbubble treatment does not impact the viability of dHL-60 cells in 4 h. **(A)** 20X representative images showing live (green) and dead (red) dHL-60 cells on a glass-bottom 96-well plate at t= 0 h and 4 h after treatment with microbubbles at 1:1, 1:2, 1:5, and 1:10 ratios or no treatment (control). Cells were stained with the LIVE/DEADTM Viability/Cytotoxicity Assay Kit at two separate time points. Scale bar, 50 μm. **(B)** Bar plots showing the viability or the percentage of live dHL-60 cells in specified conditions at t=0 h and 4 h. Each data point represents a ROI and n = 6 ROIs per condition (technical replicates). Bars show mean ± SD with the mean values written above the points. ns: p ≥ 0.05, ordinary one-way ANOVA with Tukey multiple comparisons test.

### Migration patterns of HL-60 cells and PEI microbubble conjugated HL-60 cells

The migration patterns of HL-60 cells and PEI microbubble conjugated dHL-60 cells were determined using microfluidic chemotaxis assays. We tested cells to microbubble ratios of 1:1, 1:2, 1:5, 1:10, and control with no microbubbles conjugated with cells. fMLP and LTB4 chemoattractants were used in this assay. Cells migratory behavior was different for different cell: microbubble ratios. As shown in **Figures 5A, B**, dHL-60 cells showed the highest migration for 1:1 and 1:2 ratio considering this fact that conjugating cells with microbubbles significantly decreased their migratory behavior. For 1:5 and 1:10 cell: microbubble ratio we observed the lowest migration toward both fMLP and LTB4 chemoattractants. Interestingly, PEI microbubble conjugated dHL-60 cells migration was the highest for 1:2, 1:1, 1:5 and 1:10 respectively.

**Figure 5.**
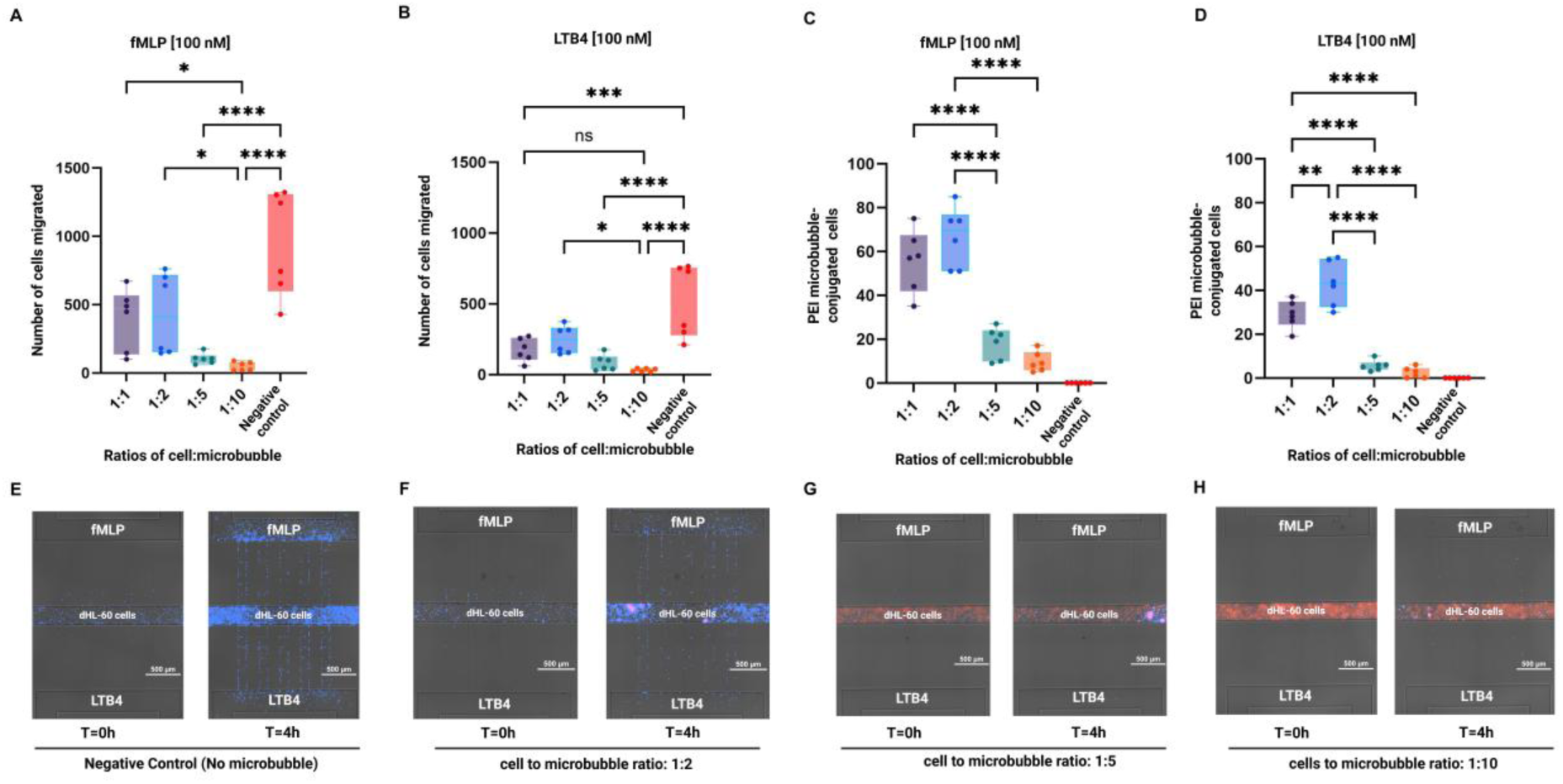
dHL-60 cells showed peak migration at a 1:2 cell-to-microbubble ratio, with reduced migration at higher ratios (1:5 and 1:10) due to impaired adhesion and mobility. **(A, B)** In comparison to the control (no microbubble conjugation), the number of dHL-60 cells migrated in response to (A) 100 nM fMLP and (B) 100 nM LTB4 at various cell-to-microbubble ratios (1:1, 1:2, 1:5, 1:10). **(C, D)** The proportion of dHL-60 cells that responded to (C) fMLP and (D) LTB4 by conjugating to PEI microbubbles at the specified ratios. **(E-H)** Representative images display the migration behavior of dHL-60 cells conjugated with varying quantities of microbubbles in response to fMLP (top chamber) and LTB4 (bottom chamber). In panel (E) dHL-60 cells (no microbubble conjugation) show strong and directed migration toward both chemoattractants, serving as a negative control. In panel (F), cells conjugated with microbubbles at a 1:2 cell-to-microbubble ratio still demonstrate effective chemotaxis. As shown in panel (G), increasing the microbubble load to a 1:5 cell-to-microbubble ratio results in a noticeable reduction in chemotactic migration. In panel (H), where the ratio reaches 1:10, dHL-60 cells completely lose their ability to migrate toward either fMLP or LTB4, suggesting that excessive microbubble conjugation severely inhibits migratory capacity. Individual data points are displayed in box plots. Data represents mean ± SD from six technical replicates (n = 6). Statistical analysis was performed using one-way ANOVA with Tukey’s post-hoc test. *p < 0.05, **p < 0.01, ****p < 0.0001; ns: not significant. Individual data points are shown as box plots.

We observed that more of the PEI microbubble conjugated HL-60 cells migrated towards the fMLP and LTB4 chemoattractant reservoir for certain ratios. We found that there was a significant difference in the number of cells that migrated towards fMLP and LTB4 when the cell to microbubble ratio increased **(Figure 5C, D). Figures 5E–H** show representative images illustrating the migration behavior of dHL-60 cells conjugated with varying amounts of microbubbles in response to fMLP (top chamber) and LTB4 (bottom chamber).

### PEI microbubbles do not independently induce migration

After identifying the migration patterns of different ratios of dHL-60: PEI microbubble conjugates towards two different chemoattractants (fMLP and LTB4), we aimed to demonstrate that the microbubbles do not induce migration independently. To test this, we conducted a chemotaxis assay using a 1:2 ratio of dHL-60: PEI microbubbles in the presence of chemoattractants and without any chemoattractant (**Figure 6A, B**).

**Figure 6.**
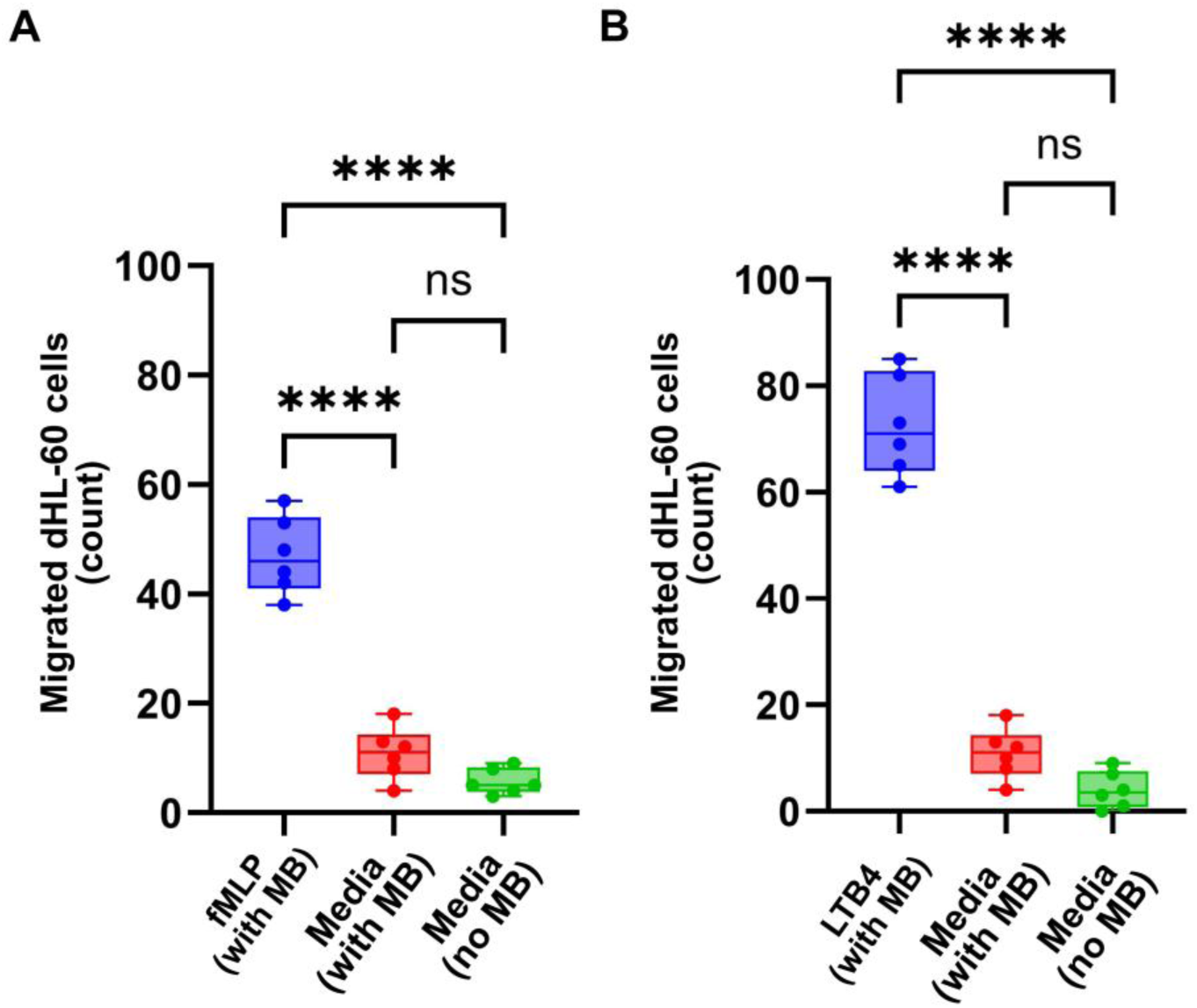
Microbubbles alone do not induce migration. Quantification of dHL-60 cell migration in response to **(A)** fMLP and **(B)** LTB4 compared to control conditions (no chemoattractant). The lack of migration in cells exposed to microbubbles in the absence of a chemoattractant (negative control) suggests that microbubbles by themselves do not promote migration. Technical replicates (n = 6) were performed for each condition. Data are presented as mean ± SD. Statistical significance was assessed using an unpaired two-tailed Student’s t-test. ****p < 0.0001.

The 1:2 dHL-60: PEI microbubble ratio without chemoattractant served as a negative control. The results showed that in the absence of chemoattractants, the conjugates exhibited no significant migration. This lack of migration in the negative control indicates that microbubbles do not independently induce. The minimal migration observed was attributed to the random movement of the conjugates within the microfluidic chip, further confirming that the directed migration was specifically due to the presence of chemoattractants. This experiment underscores that the microbubbles themselves do not have an inherent tendency to migrate.

### Increased ratio of microbubble conjugation to dHL-60 cells reduces their velocity

The velocity of dHL-60 cells in response to two distinct chemoattractants, fMLP and LTB4, under various microbubble (MB) conjugation settings (0 MB, 1 MB, and 2 MB per cell) is shown in **Figure 7A, B**. The data clearly show that, under both fMLP and LTB4 circumstances, dHL-60 cells without microbubbles (0 MB) had the highest velocities. These findings suggest that when the ratio of microbubbles to cells exceeds a certain threshold, the conjugated microbubbles impair the cells’ ability to adhere to the substrate, thereby limiting their migration toward chemotactic signals. Cells conjugated with 1 MB demonstrate a moderate drop in velocity in comparison to the 0 MB group in both chemoattractant conditions, but cells conjugated with 2 MB show the most noticeable decrease in velocity. This dose-dependent decrease in velocity suggests that adding more microbubbles further limits cellular mobility, possibly as a result of cytoskeletal dynamics being impacted by greater physical forces. Moreover, in all microbubble groups, cells exposed to fMLP exhibit somewhat higher velocities than those under LTB4 circumstances. This finding is consistent with earlier research showing that fMLP is a strong chemoattractant for neutrophil-like cells that causes robust migration reactions(50). **Figure 7A** presents the average velocity of dHL-60 cells conjugated with varying microbubble to cell ratios, while **Figure 7B** illustrates the single-cell velocity distribution across these different ratios.

**Figure 7.**
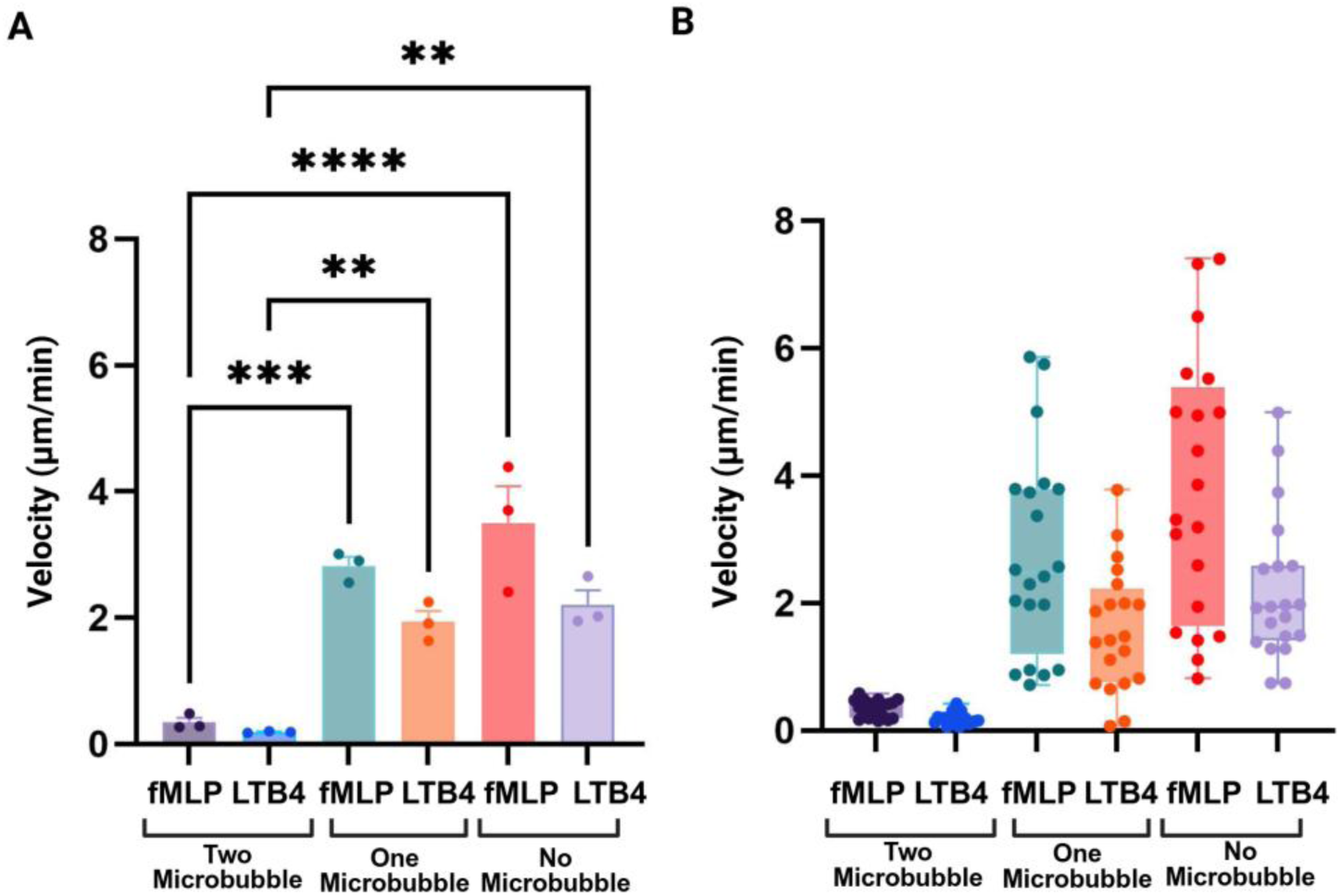
Higher microbubble binding reduces cell velocity. **(A)** The velocity of dHL-60 cells conjugated with varying quantities of microbubbles (2 MB, 1 MB, or no microbubbles) is shown in the right panel. Increased microbubble binding (2 MB) dramatically decreases motility, indicating a restricting effect on cell movement, whereas cells lacking microbubbles (0 MB) show the maximum velocity. **(B)** Average velocities under each condition confirm that increased microbubble loading correlates with decreased cell speed, with the lowest velocity observed in the 2 MB group and the highest in cells without microbubbles. Technical replicates (n = 3) were performed for each condition. Data are presented as mean ± SD. Statistical significance was assessed using one-way ANOVA. **p < 0.01, ***p < 0.001, ****p < 0.0001.

### Effect of ultrasound and microbubble conjugation on neutrophil cell migration in response to chemotactic signals

To investigate whether ultrasound exposure alters neutrophil migratory behavior, the percentage of cells migrating toward the chemoattractants fMLP and LTB4 was quantified under four ultrasound power conditions (negative control: no ultrasound, low, medium, and high power). As shown in **Figure 8**, neutrophil migration toward both fMLP and LTB4 remained comparable across all ultrasound power levels. For fMLP stimulation, no statistically significant differences were detected between control and ultrasound-treated groups. Similarly, migration toward LTB4 was consistently higher but again showed no significant differences among the different ultrasound conditions.

**Figure 8.**
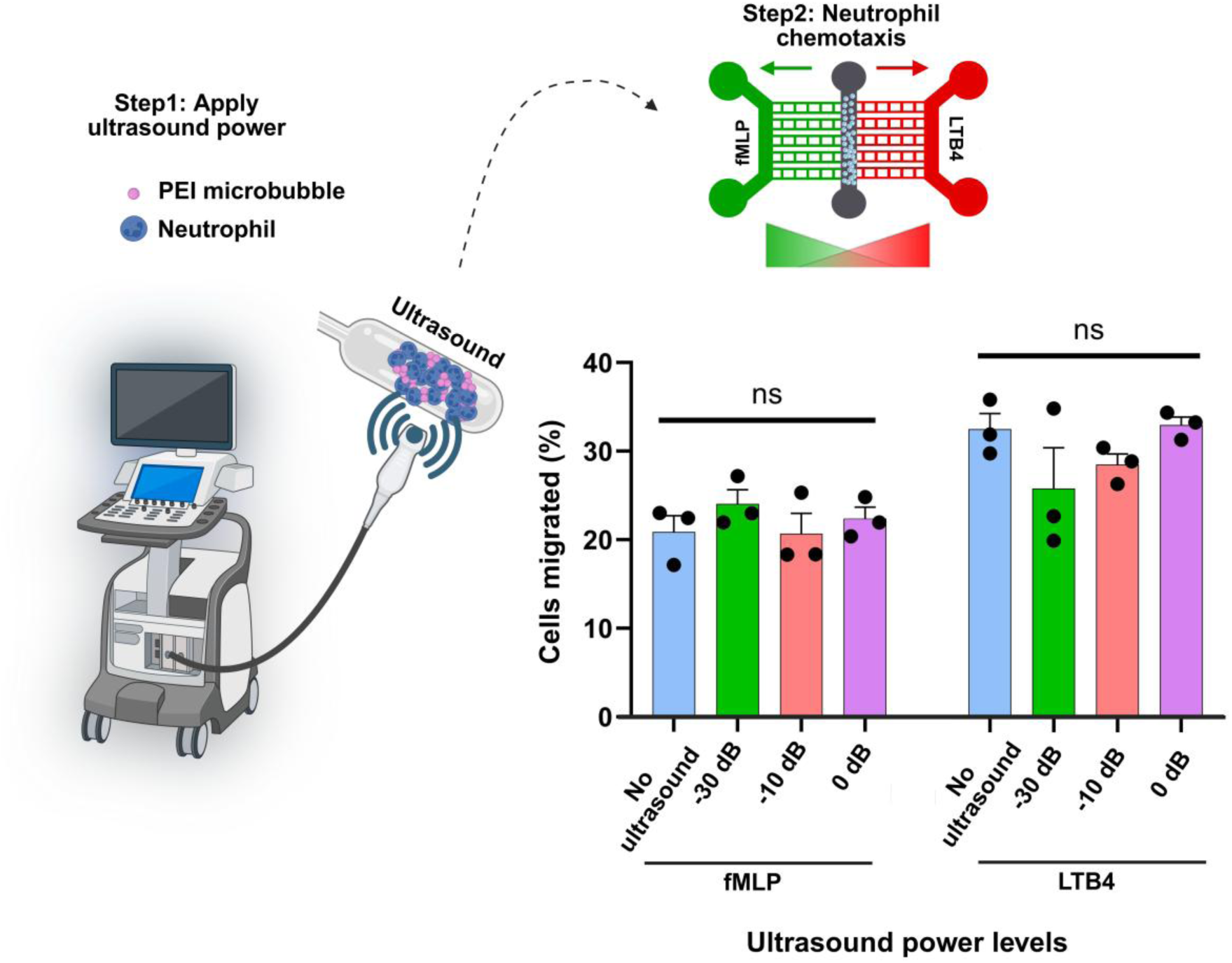
Effect of ultrasound power on neutrophil migration toward fMLP and LTB4. The percentage of migrated neutrophils was quantified under four conditions: negative control (no ultrasound), low, medium, and high ultrasound power. Each bar represents the mean ± SD of three independent experiments (black dots). No significant (ns) differences were observed among ultrasound power levels for either chemoattractant, indicating that ultrasound exposure did not alter neutrophil chemotactic activity toward fMLP or LTB4.

These results indicate that the applied ultrasound power levels did not significantly influence neutrophil chemotactic responsiveness to either fMLP or LTB4. Thus, within the tested range, ultrasound exposure appears to have minimal or no modulatory effect on neutrophil migratory capacity.

## Discussion and Conclusion

There is established research on improving immune cell infiltration into tumors using focused ultrasound and bubbles. However, this relies on increasing permeability and allowing uncontrolled immune cell infiltration(1,51). In solid tumors, where immune infiltration is frequently inadequate or spatially limited, chemotactic migration is a crucial step in the efficient utilization of immune cells for cancer therapy (52,53). The natural recruitment of immune effector cells, including neutrophils, is restricted by the complex and frequently immunosuppressive microenvironments created by tumors (54,55). Clinically, adoptive immune cell therapies such as CAR-T and NK cell infusions face major translational challenges due to limited trafficking to solid tumors, extensive ex vivo processing, and systemic toxicities. Therefore, developing a controllable, noninvasive method to guide or enhance immune cell infiltration directly in circulation could overcome a critical barrier in immuno-oncology. Our goal is to overcome this significant obstacle by employing ultrasound-guided microbubble-conjugated cells to enhance or restore directed movement by external control. Immune cells can more effectively perform cytotoxic, pro-inflammatory, or remodeling actions when they are able to locate precisely within tumor tissues because of efficient chemotaxis(52). Translationally significant, improved chemotactic guidance can decrease off-target activity in healthy tissue, increase local immune responses, and increase immune cell density at the tumor site (53,56). It is well recognized that solid tumors have immunosuppressive barriers, thick extracellular matrix, and aberrant vasculature, all of which prevent immune cells like neutrophils from infiltrating. Normal tissues usually do not have these structural and functional defects. Sonoporation can temporarily increase vascular and tissue permeability during ultrasound exposure. Because of the increased permeability and retention effect, this is more likely to occur in tumor vasculature. This action makes it easier for immune cells and microbubbles to preferentially accumulate in tumor tissue as opposed to healthy tissue. Additionally, focused ultrasound enables spatially controlled activation, which reduces off-target effects and improves the specificity of immune cell delivery to cancerous sites by enabling us to apply ultrasound only to tumor regions. When combined, this platform makes use of the physical control provided by ultrasound as well as the pathophysiological variations of tumors, making it a viable approach for targeted immunotherapy of solid tumors.

The use of microbubble conjugation and ultrasound introduces a physical and controllable approach to enhance immune cell infiltration into tumors, potentially overcoming the limitations of natural chemotaxis. According to earlier research, ultrasound can promote immune cell infiltration through improved mechanical transport, such as NK cell penetration into tumors(57). Another study by Joiner et al. have demonstrated that application of low intensity focused ultrasound with microbubbles can help activate the innate immune response for treatment of pancreatic cancer in murine models. However, the progression of disease impacted the treatment outcomes, with larger tumors being more difficult to treat due to the immunosuppressive microenvironment (58). Other studies have shown the ability to use ultrasound and microbubbles for sonopermeation, causing subsequent T-cell infiltration to the tumor(59,60). While these studies have promising results, the ability to control the type of immune cells delivered and ability to bypass the immunosuppressive microenvironment can be advantageous in improving therapeutic outcomes.

There is currently no way to selectively deliver specific immune cells to tumors by pre-conjugating them to microbubbles (MBs), which allows for immune cell-specific guidance to solid tumors. To test the feasibility of this approach, we evaluated whether PEI-coated microbubble attachment affects the viability or migration of dHL-60 cells. The LIVE/DEAD cytotoxicity assay showed no significant difference in viability between microbubble-conjugated and control cells, indicating that conjugation did not compromise cell health. Next, we characterized the migration of different cell-to-microbubble ratios under the influence of chemoattractants (fMLP and LTB4) on a microfluidic device. The purpose of performing the chemotaxis assay was to determine if the PEI microbubble-conjugated dHL-60 cells followed normal migration patterns. It was observed that the dHL-60 conjugated PEI microbubbles migrated more toward the fMLP in comparison to the LTB4. We found that a ratio of 1:2 showed the highest migration, followed by a 1:1 ratio, out of all the experimental ratios for both the chemoattractants. On the contrary, ratios of 1:5 and 1:10 inhibited the migration of PEI microbubble-conjugated dHL-60 cells due to the overloading of dHL-60 cells with the microbubbles which inhibited the migration towards both the chemoattractant. After identifying the migration pattern, we also validated if the microbubbles did not induce migration by themselves. We validated it by visualizing the migration pattern 1:2 ratio of dHL-60: PEI microbubbles in the presence and absence of chemoattractants. We observed that, in the absence of the chemoattractants, the conjugates showed no significant migration which underlines that the microbubbles themselves do not have any tendency to migrate. Lastly, we studied whether the PEI microbubbles adhered to the cells in the expected proportions. We quantified the number of PEI microbubbles (one, two, and no bubbles) and the speed of the conjugates adhering to dHL-60. It was observed that the speed of the conjugates depended on the number of microbubbles attached to the dHL-60 cells. This concluded that the more the PEI microbubbles attached to the cells lowered their migration speed towards the chemoattractants.

In addition to these experiments, we also examined the impact of ultrasound exposure on neutrophil migration behavior. Four different ultrasound power levels were tested, including a negative control with no ultrasound, low power (–30 dB), medium power (–10 dB), and high power (0 dB). The cells were exposed to these ultrasound conditions in the presence of fMLP and LTB4 gradients to evaluate potential changes in chemotactic behavior. Interestingly, we observed that ultrasound exposure across these power levels did not significantly alter the motility or directional migration of neutrophils toward either chemoattractant. These findings suggest that, within the tested range, ultrasound stimulation does not impair neutrophil migratory function, further supporting the compatibility of ultrasound-assisted delivery approaches with immune cell behavior.

Future explorations should aim to deepen our understanding of the immune response cascade initiated by these interactions. Investigating how different immune cells react to PEI microbubbles and deciphering the subsequent effects on the tumor microenvironment will provide valuable insights. These studies will facilitate the refinement of ultrasound-assisted delivery techniques, ensuring a more robust and comprehensive approach to treating solid tumors effectively.

Building upon these findings, a potential future direction involves in vivo immune cell selection and targeting. In principle, circulating immune cells could be captured via antibody-mediated binding to PEI-coated microbubbles and acoustically deposited at tumor sites. This circulation-based approach would bypass ex vivo processing steps, offering a more patient-friendly and scalable strategy for immune cell delivery.

This study presents a proof-of-concept platform demonstrating that microbubble conjugation to neutrophil-like dHL-60 cells is safe, non-cytotoxic, and compatible with essential immune cell functions. We showed that cell viability remained high (above 88%) after conjugation and that, when the microbubble-to-cell ratio is appropriately optimized, the chemotactic migration of these cells toward distinct chemoattractants (e.g., fMLP and LTB4) remains largely unaffected. To further validate the physiological relevance of our platform, we conducted additional experiments using primary human neutrophils, demonstrating that ultrasound power can effectively guide their migration without impairing motility. These results indicate that microbubble conjugation preserves critical aspects of cell health and directed migration in both model and primary immune cells, supporting the potential for immune cell delivery applications.

In conclusion, this study establishes a foundational platform for ultrasound-guided, microbubble-based immune cell delivery to tumors. By preserving cell viability and chemotactic responsiveness, this approach addresses key barriers to solid tumor immunotherapy, limited infiltration, hostile stroma, and procedural constraints, offering a scalable, patient-friendly method to potentiate existing immune therapies and improve clinical response rates. With continued refinement of the microbubble formulation and validation in in vivo tumor models, this technology has the potential to transform the way immune cells are deployed in cancer treatment.

## Conflict of Interest

The authors declare that the research was conducted in the absence of any commercial or financial relationships that could be construed as a potential conflict of interest.

## Author Contributions

HRB, CNJ and SRS supervised general work. HRB and ES optimized the microfluidic chemotaxis system. HRB and RG conducted chemotaxis experiments involving microbubbles and their injection into the microfluidic system. HRB and RG worked on optimizing the microbubble and dHL-60 cells ratio, and the injection process into the microfluidic system. HRB, SS, and APJ performed HL60 cell culture for the viability assay. GR and RG developed and optimized the microbubble generation process. HRB, APJ, and RG analyzed the chemotaxis data. SS and APJ conducted the live dead assay and analyzed the resulting data. HRB, GR, SRS, AA conducted the ultrasound section of the experiment. All authors contributed to the article and approved the submitted version. All authors assisted in writing and editing the manuscript.

## Acknowledgments and Funding

The authors would like to express their sincere gratitude to Dr. Jacques Lux, Director of the Translational Research in Ultrasound Theranostics (TRUST) Program at UT Southwestern Medical Center, and the TRUST team for their invaluable support and assistance in providing access to the ultrasound equipment and facilitating the ultrasound experiments conducted in this study. This research was supported in part by the Texas Instruments (TI) Fellowship Award from the Dean’s Office at The University of Texas at Dallas and by the National Institute of General Medical Sciences (NIGMS) under grant number 5R35GM133610-02.

## Notes

### Competing Interest Statement

The authors have declared no competing interest.

